# Single molecule imaging reveals extracellular signal-regulated kinases dependent signal responsive spatial regulation of translation in cardiomyocytes

**DOI:** 10.1101/2025.05.05.652192

**Authors:** Itai Erlich, Guy Douvdevany, Rami Haddad, Benjamin L. Prosser, Izhak Kehat

## Abstract

**Background:** Translational control of gene expression is crucial in cardiomyocytes, particularly in response to hypertrophic stimuli. The extracellular signal-regulated kinases (ERK) pathway plays a key role in inducing cardiac hypertrophy and in regulating specific protein translation. However, it is unclear how the specificity is achieved, and the spatiotemporal dynamics of protein translation remain unexplored.

**Methods:** We employed single-molecule imaging of nascent peptides (SINAPs) reporters to visualize and analyze the translation dynamics in single adult rat ventricular cardiomyocytes and tracked active translation sites at high spatiotemporal resolution. We also examined the effects of adrenergic stimulation and the role of the ERK pathway in translation localization.

**Results:** Our findings revealed that translation sites are primarily localized near Z-lines in cardiomyocytes, with some sites being highly dynamic and moving during translation. The 3’ untranslated regions did not significantly change the localization of translation. Many translation sites co-localized with microtubules, and their movement predominantly occurred along microtubular tracks. Adrenergic stimulation led to a transient shift in translation activity toward the peri-nuclear region, peaking at 12 hours and requiring ERK pathway activity for this localization change.

**Conclusions:** Our high-resolution single-cell study demonstrates that protein translation in cardiomyocytes is dynamic and responsive to hypertrophic stimuli in an ERK-dependent manner. The localized translation mechanism allows cardiomyocytes to rapidly adapt to changing environments by preferentially translating mRNAs in the peri-nuclear region.

These findings provide new insights into the spatial regulation of translation in cardiomyocytes and its role in cardiac hypertrophy.

## INTRODUCTION

Translational control of gene expression dictates the early changes in gene expression in response to hypertrophic stimuli in cardiomyocytes.^1^ We and others have shown that such hypertrophic stimuli elicit the activation of the extracellular signal-regulated kinases (ERK) pathway, which induces cardiac hypertrophy.^2,3^ ERK can regulate protein translation by inducing phosphorylation of components of the translational machinery, such as the translation factors eIF4E and 4EBP1.^4,5^ However, such activation does not globally affect protein translation, but rather appears to influence the translation of specific genes.^6^ It is not clear how this selectivity is achieved.

The local synthesis of proteins enables high concentrations of proteins where they are needed, reducing the cost of protein trafficking and facilitating proper folding while minimizing unwanted interactions.^7^ Additionally, localized translation provides spatial and temporal regulation of responses to external stimuli. For instance, through local protein synthesis, neurons can exhibit localized responses to serotonin stimulation within specific axonal branches, independent of the cell body.^8^

We previously discovered that mRNAs and ribosomes are localized in cardiomyocytes and positioned around both sides of sarcomeric Z-lines.^9^ Pulse labeling with puromycin or methionine analogs to image translation revealed localization of translation at these sites.^9,10^ However, these techniques necessitate cell fixation and fail to capture the spatiotemporal dynamics of protein translation at high resolution, which remain unexplored in cardiomyocytes.

Recent advancements in single-molecule imaging techniques have allowed visualization of nascent peptide translation dynamics using an mRNA reporter.^11–14^ The single-molecule imaging of nascent peptides (SINAPs) system employs multiple fluorescently labeled antibodies that attach to an array of epitopes added to the N-terminus of a protein open reading frame.^12^ These single-chain antibodies, fused to mature fluorescent proteins, rapidly label nascent reporter peptides as they exit the ribosome channel, enabling single-molecule imaging of translation.

Here, we employed SINAPs to unravel the spatiotemporal dynamics of translation in adult cardiomyocytes and studied their responses to hypertrophic stimuli. We found that translation sites are primarily localized near Z-lines. However, while many studies suggested that mRNAs are translationally repressed during transport^15^, in cardiomyocytes, some translation sites are highly dynamic and actively translate during transit. These movements occur predominantly along Z-line in the short axis of the cardiomyocyte, or perpendicularly to it, along the long axis of the cardiomyocyte. The movement of active translation sites along the long axis mainly occurs along microtubule tracks. We examined the role of 3’ untranslated regions (UTRs) and show they have little influence on the site of translation. Adrenergic stimulation led to the activation of protein translation in the peri-nuclear area, peaking at 12 hours and then gradually declining. Importantly, the activation of peri-nuclear protein translation was found to depend on the activity of the ERK pathway.

Our data demonstrate that localized translation in cardiomyocytes is dynamic and spatially responsive to adrenergic stimulation in an ERK-dependent manner.

## METHODS

Data supporting this study’s findings are available from the corresponding authors upon reasonable request. An expanded Materials and Methods section can be found in the Supplemental Material.

Our study examined male and female animals equally, and similar findings are reported for both sexes. All results are reported in compliance with the Animal Research Reporting of In Vivo Experiments (ARRIVE) guidelines.

During the preparation of this work, the authors used Microsoft Copilot to edit text and grammar and improve readability. The authors reviewed and edited the content as needed and take full responsibility for the publication’s content.

## RESULTS

### The single-molecule imaging of nascent peptides (SINAPs) reporter can identify translation sites in cardiomyocytes

The components of the SINAPs translation reporter system^12^ were cloned and expressed in two adenoviral vectors. The first vector contained the SINAPs reporter, which encodes 24 SunTag domains, an open reading frame of 240 amino acids, followed by an auxin-induced degron (AID) sequence. The second vector encoded a superfolder GFP attached to a single-chain variable fragment (scFV) of the GCN4 antibody that can bind to the SunTag domains of the reporter (Figure 1A). The second vector also encoded the *Oryza Sativa* transport inhibitor response 1 (OsTIR1) for degrading the C-terminus AID-containing reporter protein in the presence of auxin Indole-3-acetic acid (IAA), after completion of translation of the reporter. We first transduced isolated adult rat ventricular cardiomyocytes (ARVMs) with the vector encoding the superfolder GFP attached to scFV alone. We expected the labeling to be diffuse but found relatively faint, localized striated labeling. Co-staining with ACTN2 antibodies confirmed that these striations correspond to Z-lines (Figure 1B and 1C). The reason for this faint labeling is unclear but likely results from weak binding of the scFV to a Z-line protein, since expression of GFP alone results in diffuse labeling.^16^ We next expressed the two adenoviral vectors of the system in ARVMs in the presence of IAA. The inclusion of the SINAPs reporter system resulted in the appearance of the expected bright fluorescent spots. We added the translation inhibitor puromycin, which resulted in near complete disappearance of these bright spots, but not of the faint striated background, confirming that they are indeed active translation site (TLS) (Figure 1C, Supplemental Video 1). Quantification of the data using an automated tool in transduced ARVMs before and after adding puromycin, showed multiple TLSs at baseline but only a negligible few after puromycin (Figure 1E). We next employed fluorescence recovery after photobleaching (FRAP) on TLSs. In accordance with previous FRAP experiments of SINAPs in U2OS cells^12^, most of the recovery occurred within 200 seconds, indicate that the SINAPS reporter mRNAs undergoes constitutive translation in ARVMs (Figure 1F and 1G). Together these data show that we can reliably detect active TLSs in ARVMs.

**Figure 1.**
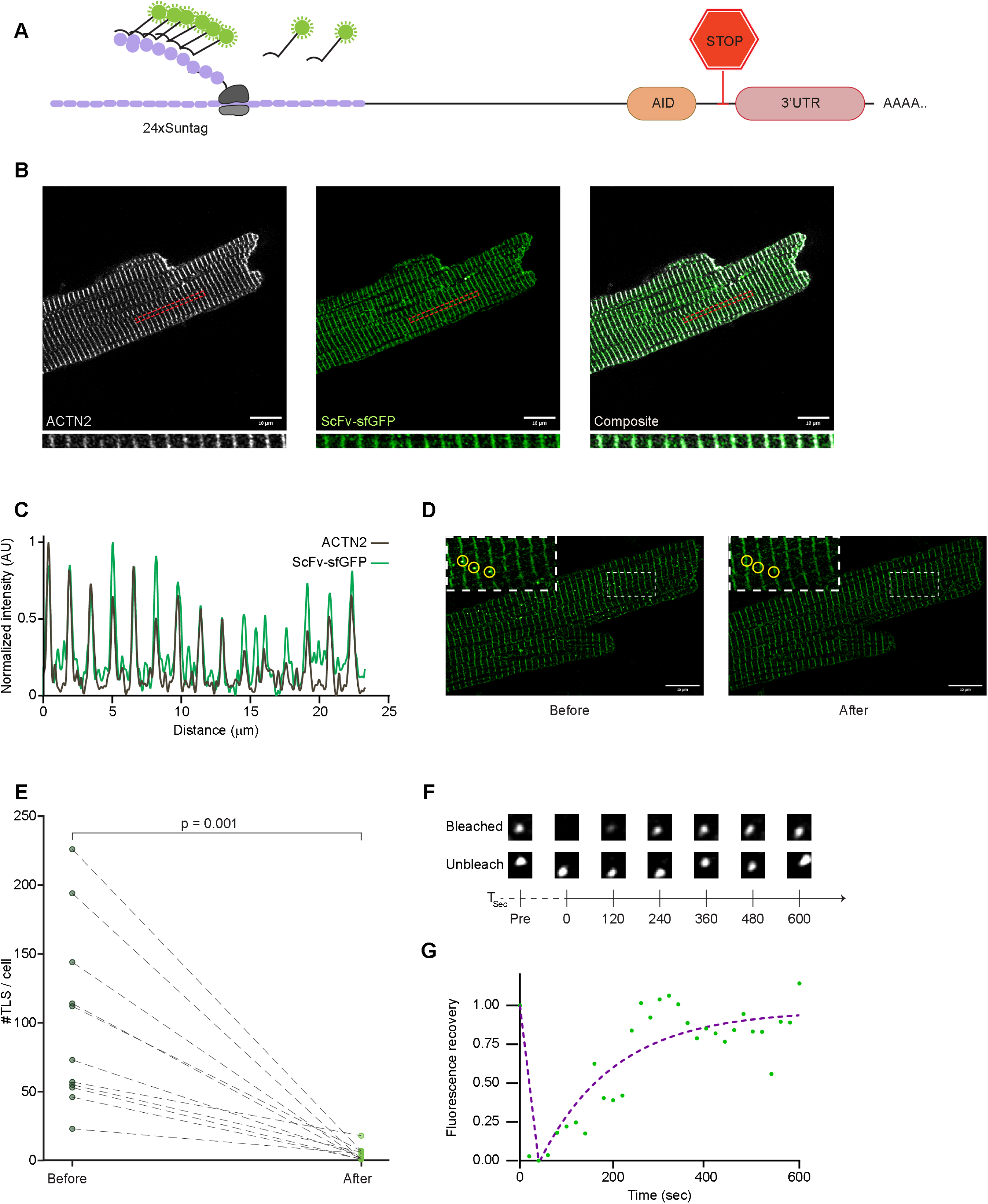
Reporter system for single-molecule imaging of nascent peptides in live cardiomyocytes. **A**, Schematic representation of the SINAPs reporter system, encoding 24 SunTag peptide domains, a 240-amino acid open reading frame, and a C-terminal auxin-induced degron (AID) sequence. Up to 24 Single chain antibodies with GFP can bind the SunTag domains upon translation, allowing the live imaging of the nascent peptide. Translation of the AID sequence results in degradation of the translated reporter, and dispersion of the GFP bound single-chain antibodies. **B**, Representative immunofluorescence image of a cardiomyocyte expressing only the single chain antibodies with suprefolder GFP (scFv-sfGFP, green) and co-stained with α-actinin antibodies (ACTN2, white) showing striated localization of scFv-sfGFP that colocalizes with Z-lines. The red dashed rectangle is shown at higher magnification at the bottom. Scale bar: 10 μm. **C**, Line scan intensity profile of the red dashed rectangle area showing co-localization of scFv-sfGFP signal (green) with ACTN2-positive Z-lines (white). **D**, Representative images of adult cardiomyocytes expressing all the components of the SINAPs reporter system showing the background cross-striated signal and the bright spots representing translation sites (yellow circles) before (left) and after (after) 3 minutes of puromycin treatment. After treatment the bright green puncta almost completely disappeared, demonstrating these were sites of active translation. Scale bar: 10 μm. **E**, The number of active translation sites (TLS) per cell was quantified before and after puromycin treatment. N=11 cells, each from an independent experiment. p = 0.001 (Wilcoxon matched-pairs signed rank test). **F**, Montage showing Fluorescence Recovery After Photobleaching (FRAP) experiments at different time points in a photobleached (upper panels) and unbleached control (lower panels) translation site. The existing nascent polypeptides (NAPs) on translation sites were photobleached at *t* = 0. Fluorescence recovery occurs as existing ribosomes continue synthesizing SunTag motifs and new ribosomes initiate translation. **G**, Representative FRAP recovery curve showing normalized fluorescence intensity over time (green dots) with fitted curve (purple line).

### Active translation sites are localized independently of the 3’ untranslated region (3’UTR)

We previously discovered that the cardiac actin (*Actc1*) mRNA was localized to Z-lines, but that the Desmoplakin (*Dsp*) mRNA was localized near the intercalated discs (ICD) in cardiomyocytes.^9^ Since the 3’ UTRs were reported to regulate localization of some mRNAs, such as the non-cardiac β-Actin (*Actb1*) mRNA^17^, we cloned and generated different SINAPs reporters encoding either no 3’UTR (null), the 3’UTR of *Actb1*, or the 3’UTR of *Actc1*. We also cloned the entire *Actc1* transcript, including its 5’ and the 3’ UTRs after the stop codon of the SINAPs to serve as a UTR, or the 3’UTR of *Dsp* (Figure S1A). We transduced ARVMs with all these different SINAPs reporters. The SINAPs reporter competes with all mRNAs for available ribosomes. As a result, fluorescent TLSs are seen only in a very small fraction of translating ribosomes in the cardiomyocyte at any given time. This sparse labeling prevents translating ribosomes from obscuring each other and allowed us to image and track single TLSs at high spatiotemporal resolution using a custom MATLAB pipeline. We observed active TLS throughout the cytoplasm with higher density near Z-lines for all the reporters (Figure 2A). The quantification confirmed the higher density of translation near Z-lines and showed that the distribution of all the reporters in respect to the Z-line was similar (Figure 2B and Figure S1B and S1C). We previously used puromycin or methionine analogs to image translation and found that in addition to the cytoplasmic Z-line localized translation, there was a high translation activity near the ICDs.^9^ We therefore specifically imaged and quantified TLSs near the edges of SINAPs transduced cardiomyocytes. As expected, this analysis showed a higher density of TLSs near the ICD, but the distribution was similar for all reporters, including the reported encoding the 3’UTR of *Dsp* (Figure 2B and 2C, and Figure S1D). Together, these data show that translation activity tends to be localized near Z-lines or close to the ICD in cardiomyocytes, but that different Actin or the *Dsp* UTRs do not significantly modify this localization.

**Figure 2.**
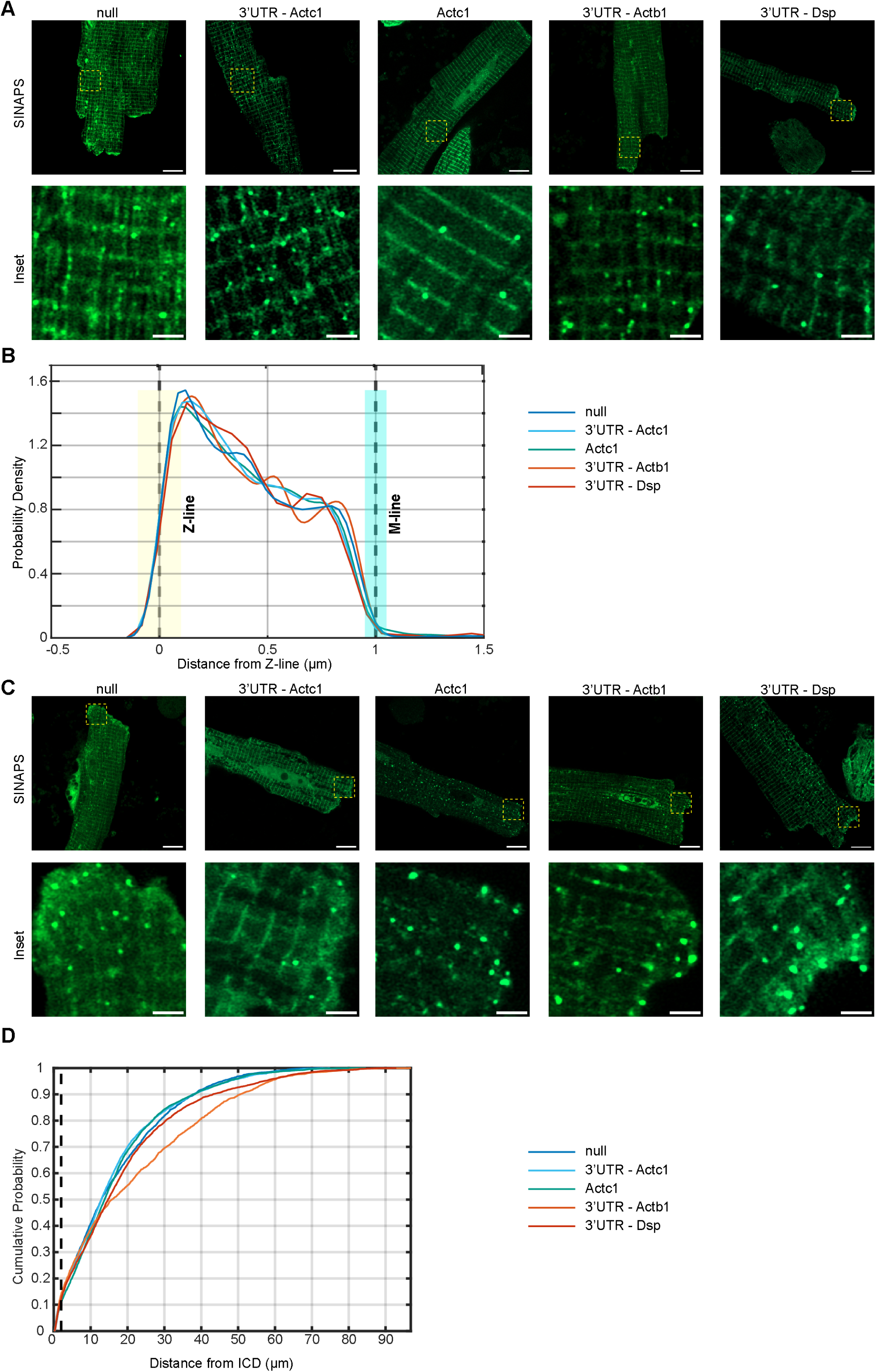
Active translation sites are localized in cardiomyocytes. **A**, Representative microscopic images of cardiomyocytes transduced with the different SINAPs reporters. Yellow dashed squares mark a cytoplasmic region which is magnified and shown in the lower panel, showing TLS distribution predominantly near Z-lines. Main image scale bar: 10 μm; inset scale bar: 2 μm. **B**, Probability density functions of TLS distances from Z-lines for different SINAPs reporters showing higher density near Z-lines for all reporters. Z-line position (0 μm) is marked by yellow dashed line and the estimated M-line position (1.0 μm) by a cyan dashed line. Semi-transparent rectangles indicate estimated Z-line (200 nm) and M-line width (100 nm). null (N=4 independent experiments, n = 22 cells, n = 2368 TLS), 3’*Actc1* (N=4 independent experiments, n = 17 cells, n = 2499 TLS), 3’*Dsp* (N=5 independent experiments, n = 25 cells, n = 2684 TLS), *Actc1* (N=6 independent experiments, n = 29 cells, n = 3513 TLS), and 3’*Actb1* (N=3 independent experiments, n = 9 cells, n = 1669 TLS). **C**, Representative microscopic images of cardiomyocytes transduced with the different SINAPs reporters. Yellow dashed squares mark a region near the intercalated disc (ICD), which is magnified and presented in the lower panel, showing higher density of TLSs. Main image scale bar: 10 μm; inset scale bar: 2 μm. **D**, Cumulative distribution of TLS distances from the ICD for the different reporters showing higher density of TLSs near the ICD for all reporters. null (N=4 independent experiments, n = 26 cells), 3’*Actc1* (N=4 independent experiments, n = 20 cells), 3’*Dsp* (N=5 independent experiments, n = 35 cells), *Actc1* (N=6 independent experiments, n = 33 cells), and 3’*Actb1* (N=3 independent experiments, n = 15 cells).

Previous studies employed a SINAPs reporter with 24 stem loops in the 3’UTR and expressed bacteriophage MS2 coat protein (MCP) fused with HaloTag, which binds these stem loops.^12^ This approach allows the simultaneous tracking of the movements of the mRNA and of active translation. We included such stem-loops in the 3’UTRs of several of our reporters (Figure S1A), and our data shows that this inclusion did not affect the localization of translation, as all our reporters showed similar spatial distribution. To image the reporter mRNA we expressed MCP-fused with HaloTag using an additional adenoviral vector, and applied Halo-JF635 fluorescent ligand. The MCP-HaloTag construct included a nuclear localization sequence (NLS) to reduce the background of the unbound construct, as was previously done.^12^ However, the expression of the NLS-MCP-fused with HaloTag alone still resulted in a strong striated cytoplasmic background signal (Figure S2A). When we co-expressed the MCP-HaloTag together with the SINAPs reporter, we could observe brighter spots that very likely represent the MCP-HaloTags bound to the multiple stem-loops in the reporter 3’UTR throughout the cytoplasm (Figure S2B). Imaging both SINAPs translation and the reporter-mRNA bound by MCP-HaloTag showed colocalization, further showing that we imaged active translation sites (Figure S2C). However, the strong background labeling of the MCP-HaloTag in cardiomyocytes hindered our ability to use this co-labeling approach reliably.

### Active translation sites are mobile in cardiomyocytes

We noticed that some TLSs were predominantly confined and stationary, while others moved during translation (Figure 3A). We used our pipeline to monitor their positions and movements (Figure 3B and 3C). We measured the average position of each TLS, which we refer to as the center position. We categorized TLS as confined if its 90th percentile of distances from the center position was smaller than 200nm. Mobile TLSs were defined as those with distances larger than 200nm (Figure 3D). We measured the distance of the center of position of each tracked TLS from the Z-line. We found that confined TLSs were significantly closer to the Z-line compared with mobile TLS (Figure 3E). These data show that translation in cardiomyocytes can be broadly divided to relatively stationary sites positioned near Z-lines, or to more mobile sites positioned further away from it.

**Figure 3.**
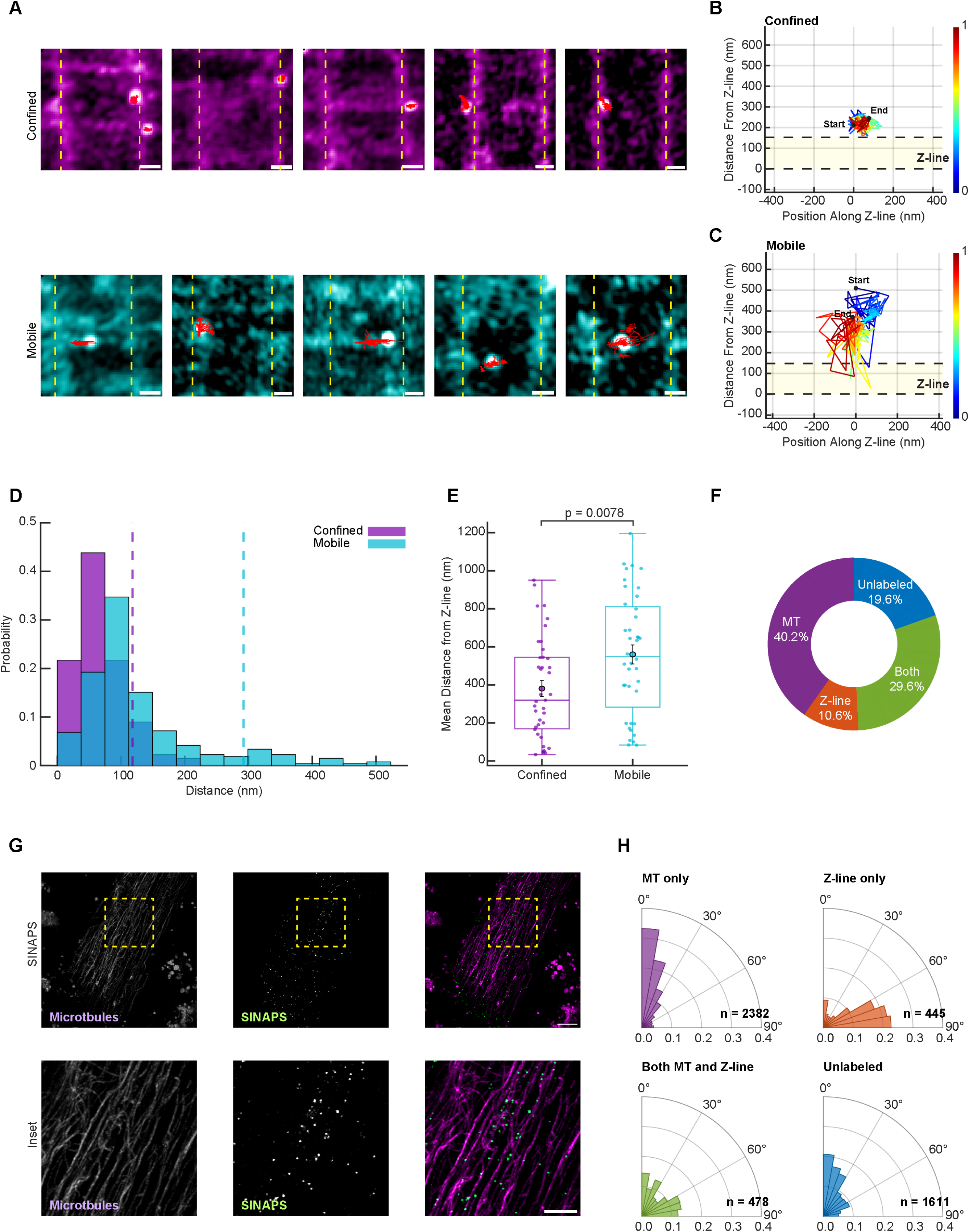
Active translation sites are dynamic in cardiomyocytes. **A**, Representative images of translation sites in cardiomyocytes exhibiting different mobility patterns. Confined translation sites (upper panels, purple) remained relatively stationary during the 5-minute live-imaging, with minimal displacement from their central position. In contrast, mobile translation sites (lower panels, cyan) displayed considerable movement. Movement trajectories are marked in red. Dashed yellow lines indicate the positions of Z-lines. Scale bar: 0.5 μm. **B**-**C**, Representative trajectory analysis of confined (B) or mobile (C) translation sites. Trajectory lines depict the displacement over the imaging interval, with colors representing time progression. Axes show the relative position of translation sites with respect to the adjacent Z-line (highlighted by yellow rectangle with black dashed borders). **D**, Representative displacement distribution histograms comparing confined (purple) and mobile (cyan) translation sites, with 90th percentile thresholds indicated by dashed lines (confined = 118.1 nm; mobile = 292.1 nm). Confined sites exhibit a narrower distribution concentrating near their central position, while mobile sites show broader displacement patterns. **E**, Box plot with individual data points showing the center positions of confined (purple) translation sites were significantly closer to Z-lines compared with that of mobile (cyan) translation sites (n= 39,40 translation sites from N = 4 independent experiments and N= 10 cells. p = 0.0078, two-sample t-test. **F**, Doughnut chart quantifying the percentage of active translation sites that were colocalized with microtubules (MT), Z-lines, both, or neither (N = 3 independent experiments, n= 123 sites and N= 10 cells). **G**, Representative images of cardiomyocytes transduced with SINAPs reporter (green) and co-stained with live-marker Tubulin Tracker (purple). Inset below at higher magnification showing many active translation sites were co-localized with microtubules. Scale bar: 10 μm, Inset scale bar: 5 μm. **H**, Translation sites trajectories with >200 nm displacement were classified by their association with microtubules (MT) or Z-lines and movement angle was plotted on polar histograms with 0° representing the long and 90° representing the short axis of the cardiomyocyte. n of trajectories is noted on each plot (N = 10 cells).

Ablation of the microtubular network was shown to result in the loss of the sarcomeric localization of mRNA and ribosome, and to their collapse to a peri-nuclear localization.^9,18^ We therefore stained live ARVMs with a fluorescent Tubulin Tracker dye and imaged TLSs and microtubules together. We quantified the co-localization of TLSs with Z-lines and with microtubules and found that 40.2% of TLSs were colocalized with microtubules. An additional 29.6% were found at junction sites and were colocalized with both microtubules and Z-lines. Some TLSs were co-localized with Z-line alone, and the co-localization of 19.6% of TLSs could not be reliably defined (Figure 3F and 3G). Together these data show that 69.8% of TLSs we imaged were colocalized with microtubules, while 40.2% were colocalized with Z-lines. To further verify that TLSs were associated with microtubules and not only colocalized with them we disrupted the network using nocodazole. In accordance with previous observations^18^, the disruption of the microtubular network resulted in loss of localization and the collapse of active translation to a peri-nuclear region (Figure S3A and S3B). We next tracked the movements of TLSs and defined the long axis of the cardiomyocyte as 0^0^ direction, and the perpendicular short axis as 90^0^. We found that TLSs that were colocalized with microtubules predominantly moved at 0^0^ direction along the direction of most microtubular tracks. In contrast, TLSs which were colocalized only with Z-lines, predominantly moved in the 90^0^ direction, in parallel to Z-lines, and TLSs associated with both moved either at 0^0^ or at 90^0^ direction, but rarely diagonally (Figure 3H, Supplemental Video 2). TLSs which were not reliably colocalized with either the Z-lines or with microtubules also predominantly moved at 0^0^ direction. It is possible that the latter are associated with other cytoskeletal elements that are aligned in the long axis such as the non-sarcomeric actin network. These data show that while translation is localized in cardiomyocytes, it is much more dynamic than previously assumed.

### Adrenergic stimulation results in a transient translation shift in cardiomyocytes

We wondered if adrenergic stimulation, a strong pro-hypertrophic signal, would affect the spatial dynamics of translation. We therefore treated ARVMs with the α-adrenergic agonist phenylephrine (PE) or vehicle and imaged them at 3,12,24, or 72 hours. We found that stimulation with PE resulted in a shift of translation activity from distal cytoplasmic sites, to area closer to the nucleus. This shift peaked at 12 hours, and gradually declined, reaching vehicle levels at 72 hours (Figure 4A). We quantified the finding by plotting the cumulative probability of finding TLSs as a function of the distance from the nucleus (Figure 4B) or compared the cumulative probability of TLSs positions withing 5 µm of the nucleus edge (Figure 4C) and found a significant shift of active TLSs location closer to the nucleus with PE stimulation. We next wanted to verify that this shift occurs in the same cardiomyocytes, and therefore imaged ARVMs with time-lapse microscopy for 6 hours after treatment with vehicle or PE. A similar shift in translation was observed, with TLSs concentrating near the nucleus starting two hours after stimulation and becoming more pronounced at six hours. No shift was observed in vehicle-treated cardiomyocytes (Figure 4D and 4E). We similarly quantified these findings and plotted the cumulative probability of finding TLSs as a function of the distance from the nucleus edge (Figure 4F) and the normalized cumulative probability of finding a TLS within 5 µm of the nucleus (Figure 4G), noting significant shift of active translation to peri-nuclear region at 2,3 and 6 hours in PE compared with vehicle treated ARVMs.

**Figure 4.**
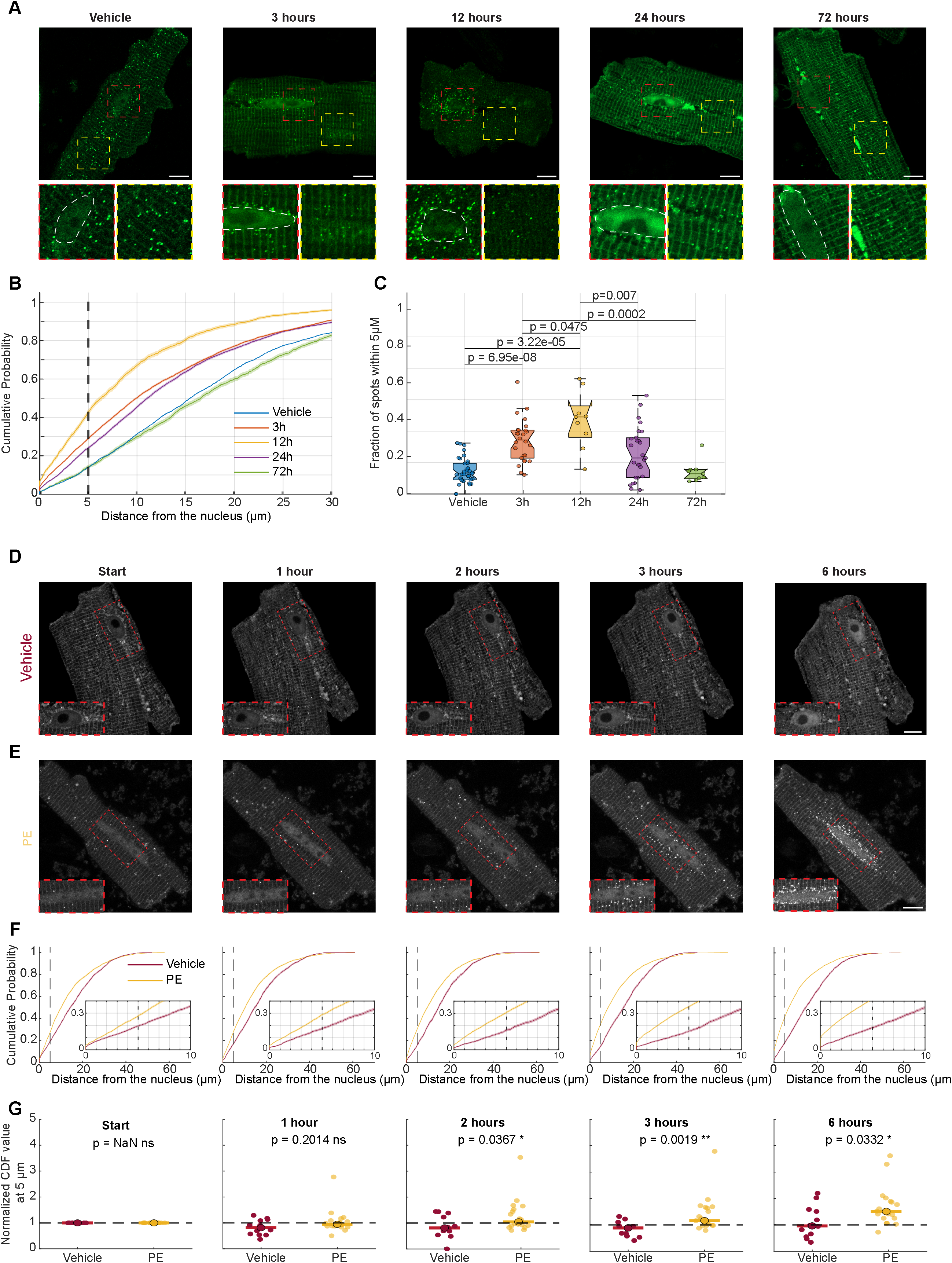
α-adrenergic stimulation results in peri-nuclear activation of translation. **A**, Representative images of transduced adult cardiomyocytes exposed to phenylephrine (PE, 50 μM) for varying durations (0, 3, 12, 24, and 72 hours). Perinuclear accumulation of active translation sites is evident after 3-12 hours of stimulation, diminishes after 24 hours, and returns to baseline distribution by 72 hours. The effect is highlighted in magnified views of the perinuclear region (red dashed squares) compared to a peripheral cytoplasmic region (yellow dashed squares). Nuclei borders are marked in white dashed line. Scale bar: 10 μm. **B**, Cumulative distribution of active translation sites positions as a function of distance from the nuclear edge. Dashed line shows 5μm distance used for further analysis. **C**, Analysis of cumulative probability values at 5 μm distance from the nucleus for each experimental condition showing peri-nuclear accumulation of active translation sites (Vehicle: N=2 independent experiments, n=36 cells; PE 3 hours: N=5 independent experiments, n=27 cells; PE 12 hours: N=2 independent experiments, n=9 cells; PE 24 hours: N=3 independent experiments, n=27 cells; PE 72 hours: N=2 independent experiments, n=9 cells). Statistical significance was determined using Benjamini-Hochberg false discovery rate (FDR) correction applied for multiple comparisons. **D-E**, Representative time-lapse images of a single adult cardiomyocyte treated with vehicle (D) or phenylephrine (PE) (E) at the starting point and after 1, 2, 3, and 6 hours. Red dashed boxes show magnified views of the perinuclear region, demonstrating accumulation of active translation sites around the nucleus only in PE treated cells. Scale bar: 10 μm. **F**, Cumulative distribution of active translation sites positions as a function of distance from the nuclear edge (vehicle – red, phenylephrine - yellow). Inset magnifies the distribution within 10 μm of the nucleus, highlighting differences between treatment groups over time. **G**, Analysis of cumulative probability values at 5 μm distance from the nucleus for each experimental condition and time-point (vehicle: N=3 independent experiments, n=12 cells; phenylephrine: N=3 independent experiments, n=19 cells). Statistical significance was determined using Mann-Whitney U test.

We wanted to examine if the shift in translation localization was specific to α-adrenergic stimulation and therefore also treated cardiomyocytes with the β-adrenergic agonist isoprenaline (ISO). Stimulation with ISO similarly resulted in a significant shift of translation to the perinuclear region (Figure S4A through S4C). To determine if the 3’UTR played a role in the response to PE we imaged ARVMs transduced with the *Dsp* 3′UTR reporter and found it similarly responded to ISO stimulation with a shift toward the nucleus (Figure S4D through S4F).

### Adrenergic induced translation shift in cardiomyocytes requires ERK activity

An adrenergic-induced shift in the localization of translation to the peri-nuclear region may be due to alterations in mRNA localization, changes in ribosome positioning, or specific activation of translation in the peri-nuclear region. To examine these possibilities we first performed single-molecule mRNA fluorescent in situ hybridization (smFISH) for 18S ribosomal RNA to detect ribosomes, as we have previously done^10^, in ARVMs treated with PE or vehicle for 6 hours. This staining did not show any collapse of ribosomes to the peri-nuclear region (Figure 5A). Ribosomes were localized in a striated pattern in the cytoplasm, with denser localization of ribosomes in the peri-nuclear region in both vehicle and PE treated cells. We next performed a similar analysis using smFISH for polyA tails to label all mRNAs, and did not observe a marked global shift in mRNA localization with PE stimulation (Figure 5B). PolyA mRNAs were localized in a striated pattern in the cytoplasm, with denser localization inside the nucleus and around it in both vehicle and PE treated cells.

**Figure 5.**
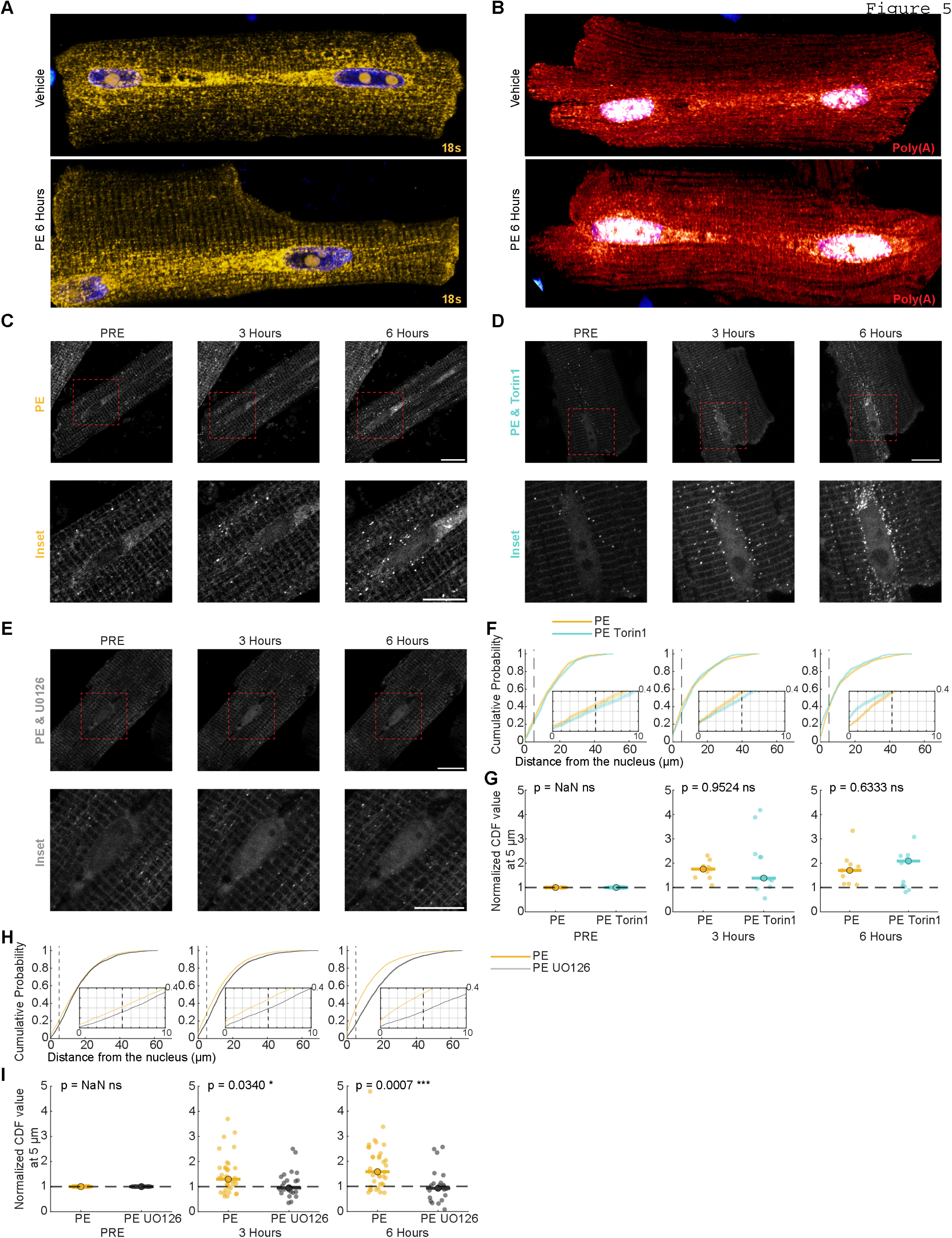
ERK activity is required for adrenergic-induced spatial control of translation. **A**-**B**, Representative images of single-molecule fluorescence in situ hybridization (smFISH) detecting 18S rRNA (A, yellow) or polyA mRNA (B, red) in adult cardiomyocytes treated with vehicle (upper panel) or phenylephrine (PE, lower panel) for 6 hours. Images do not show a major redistribution of ribosomes or mRNA in cardiomyocytes following phenylephrine treatment. **C-E**, Representative time-lapse images of transduced adult cardiomyocytes treated with phenylephrine at 0, 3, and 6 hours with addition of vehicle (C), Torin1 (D), or UO126 (E). Magnified views of the perinuclear region (red dashed squares) are shown below. Perinuclear accumulation of active translation sites is evident after 3-6 hours of stimulation in phenylephrine or phenylephrine + Torin1 treated cells, and the effect is blocked by UO126. The effect is highlighted in. Scale bar: 10 μm. **F-G**, Cumulative distribution of active translation sites and analysis of cumulative probability values at 5 μm distance from the nucleus for phenylephrine and vehicle (yellow) and phenylephrine with Torin1 (cyan). Inset magnifies the distribution within 10 μm of the nucleus, showing no significant differences between the two treatment conditions. Phenylephrine: N=3 independent experiments, n=9 cells; phenylephrine and Torin1: N=3 independent experiments, n=15 cells). Statistical significance was determined using Mann-Whitney U test. **H-I**, Same analysis as in F-G comparing phenylephrine and vehicle (yellow) and phenylephrine with U0126 (cyan). (phenylephrine: N=6 independent experiments, n=36 cells; phenylephrine and U0126: N=6 independent experiments, n=24 cells). Statistical significance was determined using Mann-Whitney U test.

The mammalian target of rapamycin (mTOR) protein kinase regulates translation and has been found to control localized translation in neurons. This effect could be inhibited by Torin1.^19^ We therefore examined translation in the presence of PE and Torin1 or PE and vehicle. The blockade of mTOR activity did not blunt the translation shift, and TLSs still concentrated in the peri-nuclear region after PE stimulation in the presence of Torin1 (Figure 5C and 5D). ERK kinases is another pathway that controls several proteins involved in translation regulation.^6^ To examine its role we repeated the PE stimulation with or without the MEK inhibitor UO126. The blockade of the ERK pathway resulted in a marked effect, and we no longer observed the shift of translation to the peri-nuclear region (Figure 5E). The quantification of the data confirmed that inhibition of mTOR activity by Torin1 did not result in significant blunting of the PE induced shift in translation towards the nucleus (Figure 5F and 5G). In contrast, the inhibition of the MEK-ERK pathway significantly blunted this shift (Figure 5H and 5I).

## DISCUSSION

Our previous studies demonstrated that the translation machinery and protein synthesis are localized on both sides of the Z-lines in cardiomyocytes.^9,10^ These observations were subsequently corroborated in skeletal muscles.^20^ These analyses, conducted on fixed cells, offered insights into the localization of translation in myocytes, but implied that protein translation was confined. Using live imaging of translation, we show here that while some translation sites reside predominantly near Z-lines and have limited motion, others are dynamic and move during the translation process. These movements do not suggest passive diffusion as they are not random and predominantly occur either along the Z-lines, or perpendicularly to them, along the long axis of the cardiomyocyte.

Many active translation sites were colocalized with microtubules, and we show here that these active sites move bidirectionally along the microtubules in the long axis of cardiomyocytes. We and others have previously shown that disruption of the microtubular network resulted in a mis-localization and collapse of mRNA and ribosomes around the nucleus.^9,18,20^ The data shown here indicates that microtubules are not only used for transport of mRNAs and ribosomes from the nucleus to their peripheral destinations, but also participate in moving actively translating ribosomes. The position of ribosomes around both sides of Z-lines allow proximity to the myofibrils but also to organelles such as the endoplasmic reticulum and mitochondria, which surround myofibrils. The role of active translation sites movements is not clear, but we speculate that it may allow nascent proteins opportunity to engage with sarcomeric structures or the surrounding organelles. A signal peptide or a specific peptide motif in the N-terminus of the nascent chain could then anchor the translation and the emerging peptide to the appropriate site.

It has long been recognized that cardiac hypertrophy is associated with increased protein synthesis, and that preferential synthesis of new ribosomes occurs within hours of hypertrophic stimuli.^21,22^ A later detailed ribosome-sequencing study showed that subsets of mRNAs are regulated at the translational level early after the induction of cardiac hypertrophy.^1^ The increase in translation of specific subsets of mRNAs suggests a mechanism that could target specific mRNAs, but such a mechanism remained elusive. Our data show that hypertrophic stimulation can alter the spatial translation landscape of cardiomyocytes by increasing protein translation around the nucleus. This mechanism can allow cardiomyocytes to preferentially translate mRNAs that are enriched in this area. For example, it was shown that β1 adrenergic receptor mRNA^23^ and Cx43 mRNA^24^ are localized in the perinuclear region in cardiomyocytes, and it is likely that many other mRNAs are localized there as well. As newly synthesized mRNAs exit the nucleus and move to the periphery, initiating translation near the nucleus can prioritize translation of these nascent mRNAs over older ones further away, coupling translation with transcription.

Many pro-hypertrophic endocrine and paracrine signals, including adrenergic signals, induce activation and phosphorylation of the ERK cascade.^25,26^ This activation results in ERK nuclear translocation in cardiomyocytes, and data suggest that this nuclear translocation is required for concentric hypertrophic growth.^27^ Very recent work, conducted independently, found that phenylephrine induces an mTORC1-independent, but an ERK-dependent pathway that drives phosphorylation of 4EBP1 at Ser64 in cardiomyocytes.^28^ Phosphorylation of 4EBP1 can decrease its binding to eIF4E and increase translation.^29^ Our data also shows a phenylephrine induced ERK-dependent and mTORC1-independent pathway. Here we show that this pathway induced activation of translation specifically in the peri-nuclear region of cardiomyocytes.

Previous use of a similar translation reporter system in other cell types yielded data that was consistent with translation parameters measured using other approaches^12,30^, but the reporter system is not without limitations. We imaged the translation of a synthetic mRNA reporter that does not encode an endogenous protein. While we showed that the 3’UTR does not play a major role in localized translation in cardiomyocytes, we cannot rule out that some specific endogenous mRNAs could exhibit different behaviors, or that translation of specific motifs could alter the localization. We also used cultured ARVMs, because high resolution imaging is not possible in vivo. While our imaging of the reporter mRNA showed it was present throughout the cytoplasm, we cannot rule out the possibility that certain subsets of ribosomes were more available and more likely to translate the reporter. We therefore cannot rule out the possibility that our assessment of the percentage of translating ribosomes associated with Z-line or microtubules could be somewhat biased.

In summary, we showed that protein translation is localized and dynamic in cardiomyocytes and involves active movement of translation sites along microtubules and Z-lines. We demonstrate that localization is largely independent of the 3’UTR. Importantly, we uncovered a signal-responsive localized translation response, which is mediated by the ERK cascade, that allows cardiomyocytes to differentially translate proteins in the peri-nuclear region in response to pro-hypertrophic adrenergic stimuli. This response can allow cardiomyocytes to translationally control the expression of specific genes and to rapidly respond to changing environments and stimuli.

## Acknowledgements

The authors wish to thank the Biomedical Core Facility at the Faculty of Medicine, Technion and the Pre – Clinical Research Authority at the Technion.

---

**Figure S1.**
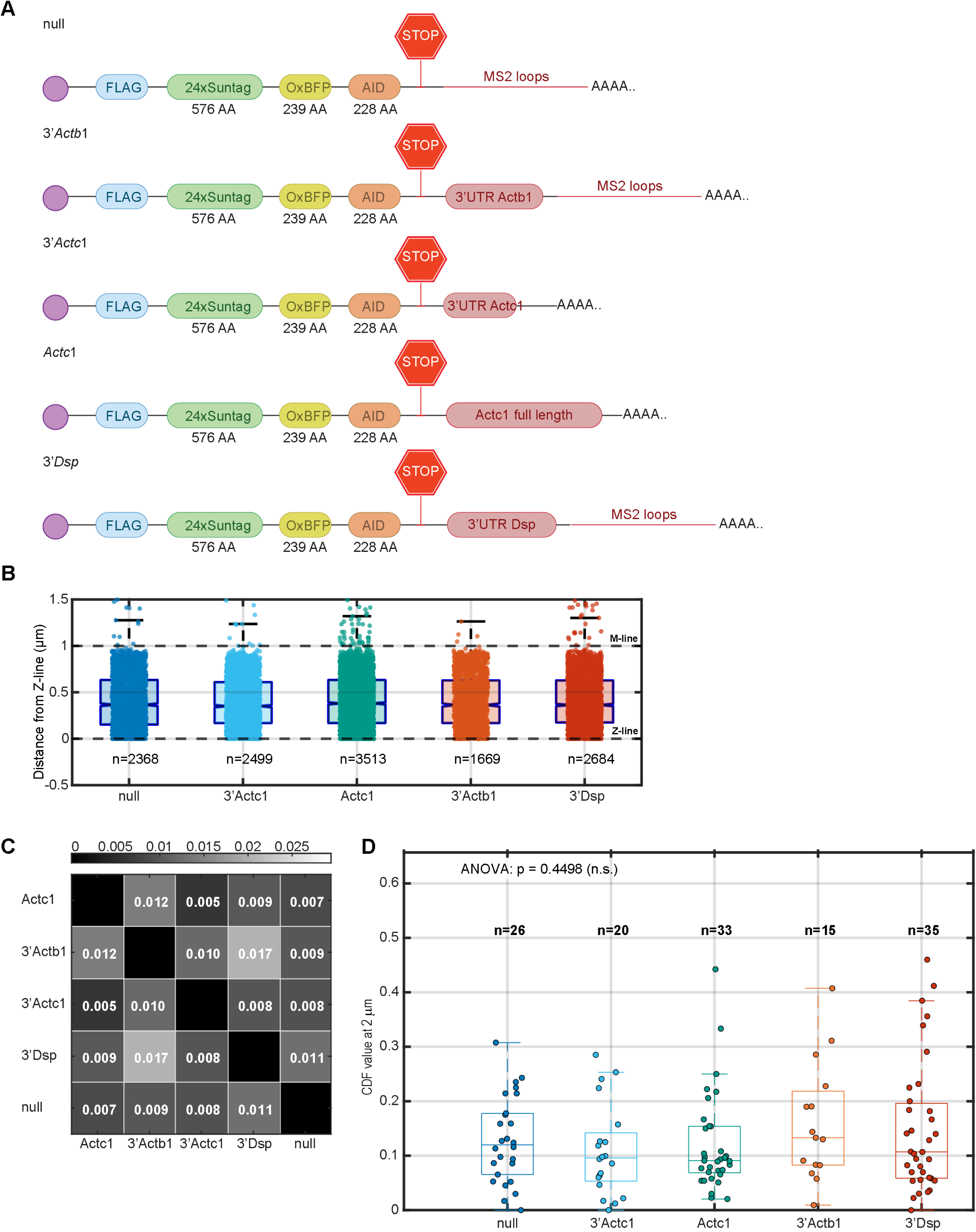
Translation localization is independent of 3’UTRs. **A**, illustration of the different SINAPs reporters used, encoding either no 3’UTR (null), the 3’UTR of *Actb1*, the 3’UTR of *Actc*1, the entire *Actc*1 transcript as UTR, or the 3’UTR of *Dsp*. The null, 3’UTR of *Actb1*, and *Dsp* reporter also encode MS2 loops. **B**, Box plots show the distribution of distances from Z-lines for TLS for the different reporter constructs. Horizontal dashed lines indicate Z-line (0 μm) and M-line (1.0 μm) positions. Mean distances from Z-lines were (mean ± SD): null (0.41 ± 0.29 μm), 3’*Actc1* (0.40 ± 0.29 μm), 3’*Dsp* (0.42 ± 0.36 μm), *Actc1* (0.41 ± 0.29 μm), and 3’*Actb1* (0.41 ± 0.28 μm). For each reporter: null (N=4 independent experiments, n = 22 cells, n = 2368 TLS), 3’*Actc1* (N=4, n = 17 cells, n = 2499 TLS), 3’*Dsp* (N=5, n = 25 cells, n = 2684 TLS), *Actc1* (N=6, n = 29 cells, n = 3513 TLS), and 3’*Actb1* (N=3, n = 9 cells, n = 1669 TLS). Statistical analysis using Kruskal-Wallis test (p=0.77) and pairwise comparisons with Mann-Whitney U tests revealed no significant differences between any reporter constructs. **C**, Heatmap displaying Jensen-Shannon (JS) divergence values between TLS spatial distributions for the different SINAPs reporters. JS divergence quantifies the dissimilarity between probability distributions on a scale from 0 (identical distributions) to 1 (maximally different). Values in each cell represent the JS divergence between the corresponding construct pairs, and all measured divergence values were below 0.02, indicating highly similar spatial distribution patterns across all tested constructs. Permutation test revealed no statistically significant differences between any reporter pairs (p>0.05), suggesting that the spatial organization of TLS events relative to sarcomeric Z-lines is highly independent of the 3’UTR. **D**, Comparison of TLS localization at 2 μm from the ICD region across SINAPs reporters. Box plots display the distribution of cumulative distribution function (CDF) values at 2 μm for each reporter, with individual data points representing measurements from single cells. For each reporter: null (N=4 independent experiments, n = 26 cells), 3’*Actc1* (N=4 independent experiments, n = 20 cells), 3’*Dsp* (N=5 independent experiments, n = 35 cells), *Actc1* (N=6 independent experiments, n = 33 cells), and 3’*Actb1* (N=3 independent experiments, n = 15 cells). Statistical significance between groups was determined by one-way ANOVA (p=0.45) followed by Mann-Whitney U tests showing no significant differences between any reporter constructs

**Figure S2.**
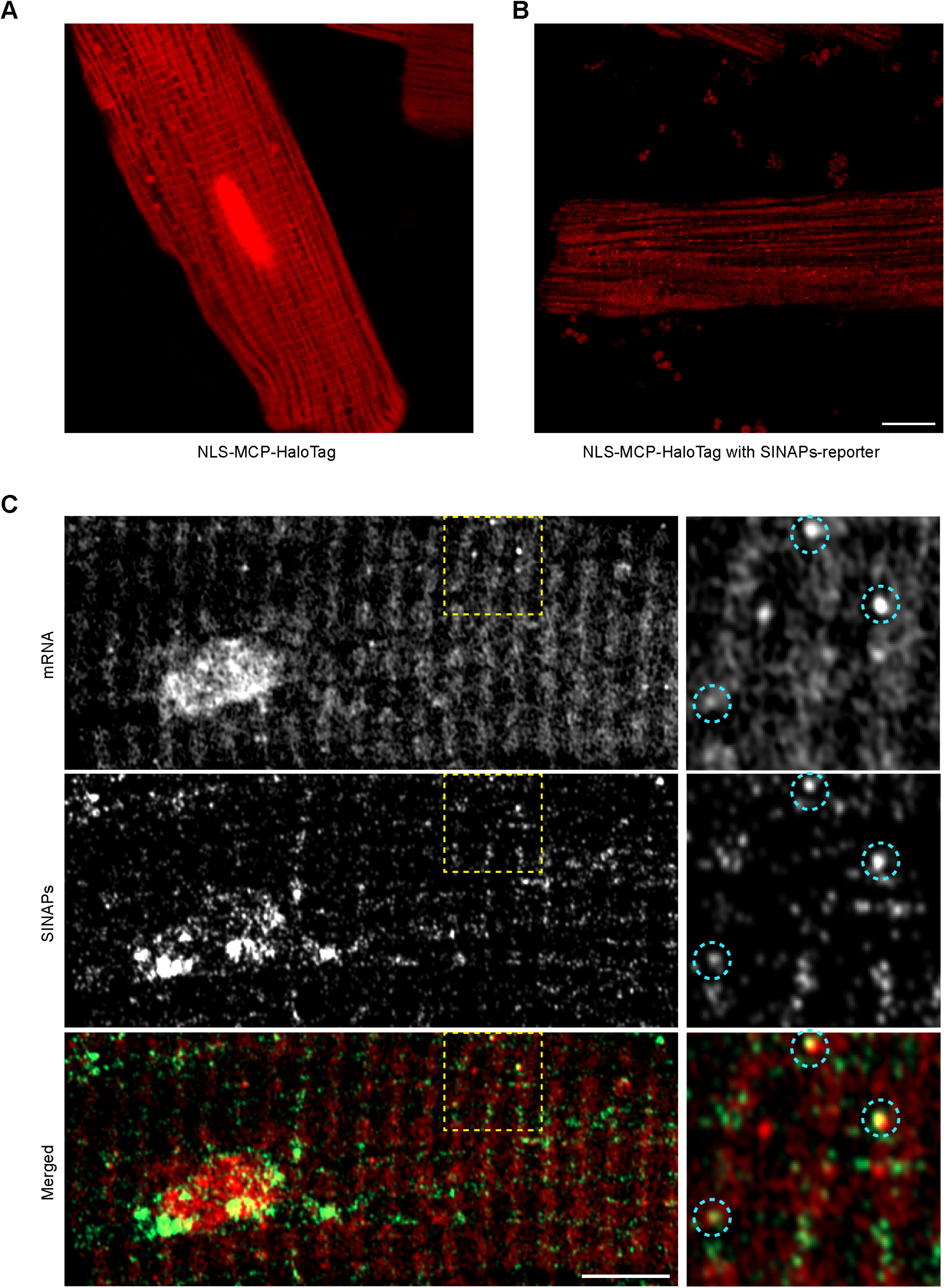
Imaging of the SINAPs reporter mRNA. **A**, Representative image of cardiomyocyte transduced with MCP-HaloTag with a nuclear localization sequence (NLS) alone and incubated with Janelia Fluor 635 Halo ligand. The image shows a strong, non-specific background cross-striated signal and a pronounced nuclear labeling. Scale bar: 10 μm. **B**, Representative image of cardiomyocytes co-transduced with MCP-HaloTag and the SINAPs reporter containing 24xMS2 loops in the 3’UTR. The specific binding of MCP-HaloTag fusion protein to MS2 loops allows a somewhat improved signal-to-noise ratio and visualization of the reporter mRNA molecules upon addition of Janelia Fluor 635 ligand. Scale bar: 10 μm. **C**, Representative images of active mRNA translation in cardiomyocytes using dual-reporter system, with imaging of reporter mRNA upon addition of Janelia Fluor 635 ligand (white in the upper panel, red in the lower merged panel), and of the SINAPs reporter (white in the middle panel, green in the lower merged panel). Actively translating mRNAs appear as yellow spots where both signals colocalize. Higher magnification of the dashed boxed region shows this localization (circles). Scale bar: 5 μm; inset scale bar: 2.5 μm

**Figure S3.**
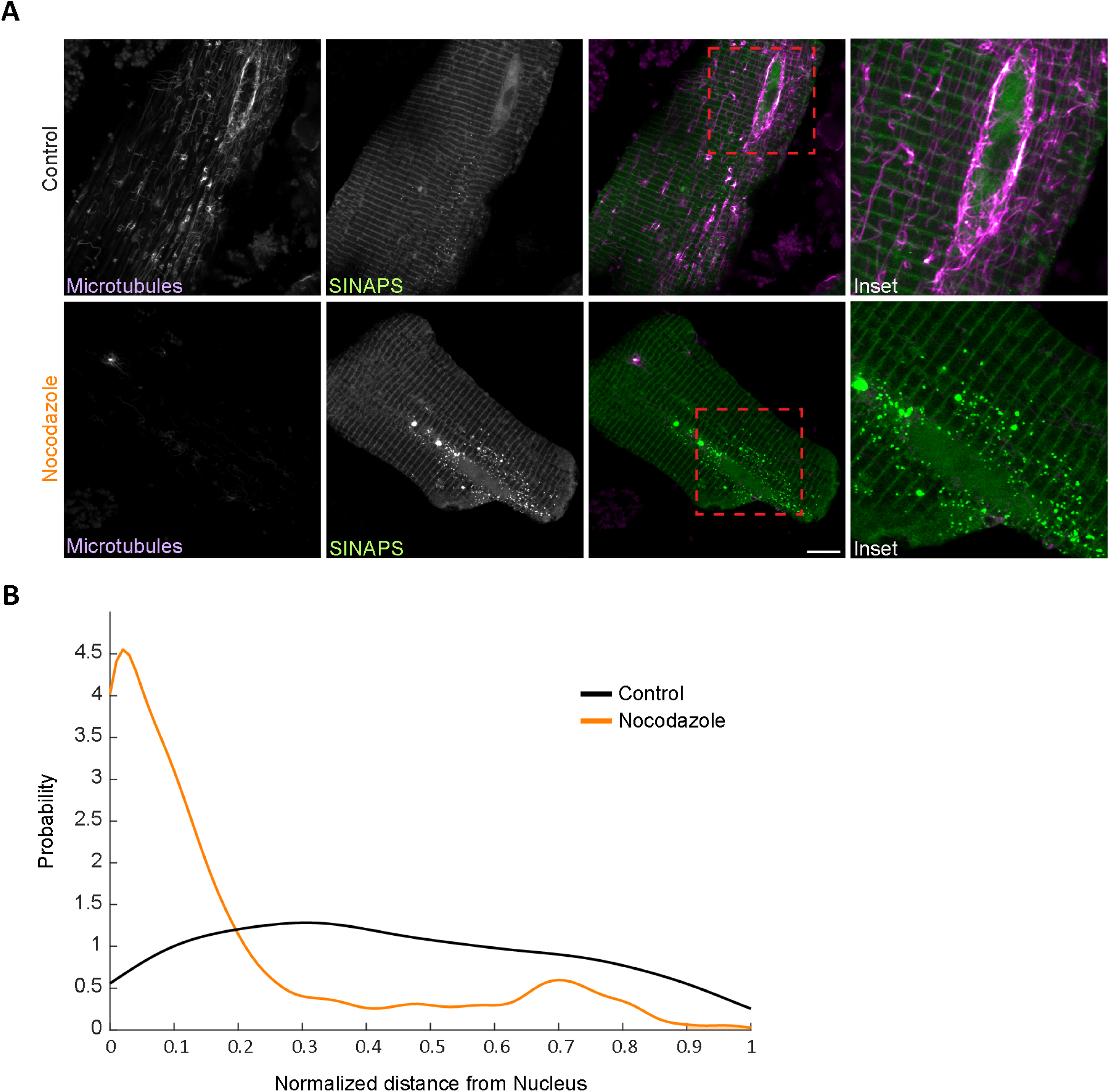
Microtubule ablation results in mislocalization and peri-nuclear collapse of translation sites. **A**, Representative images of cardiomyocytes expressing the SINAPs reporter (green) and stained for microtubules (purple) treated with control (upper) or nocodazole (10 µM) for 24 hours (lower panels). Dashed area is shown at higher magnification. Nocodazole treatment results in depolymerization of the microtubular network and perinuclear accumulation of active translation sites. Scale bar: 10 µm. **B**, Probability distribution of translation site position as a function of distance from the nuclear edge in control (black) and nocodazole-treated (orange) cardiomyocytes. N = 2 independent experiments, control n = 6 cells; nocodazole n = 7 cells).

**Figure S4.**
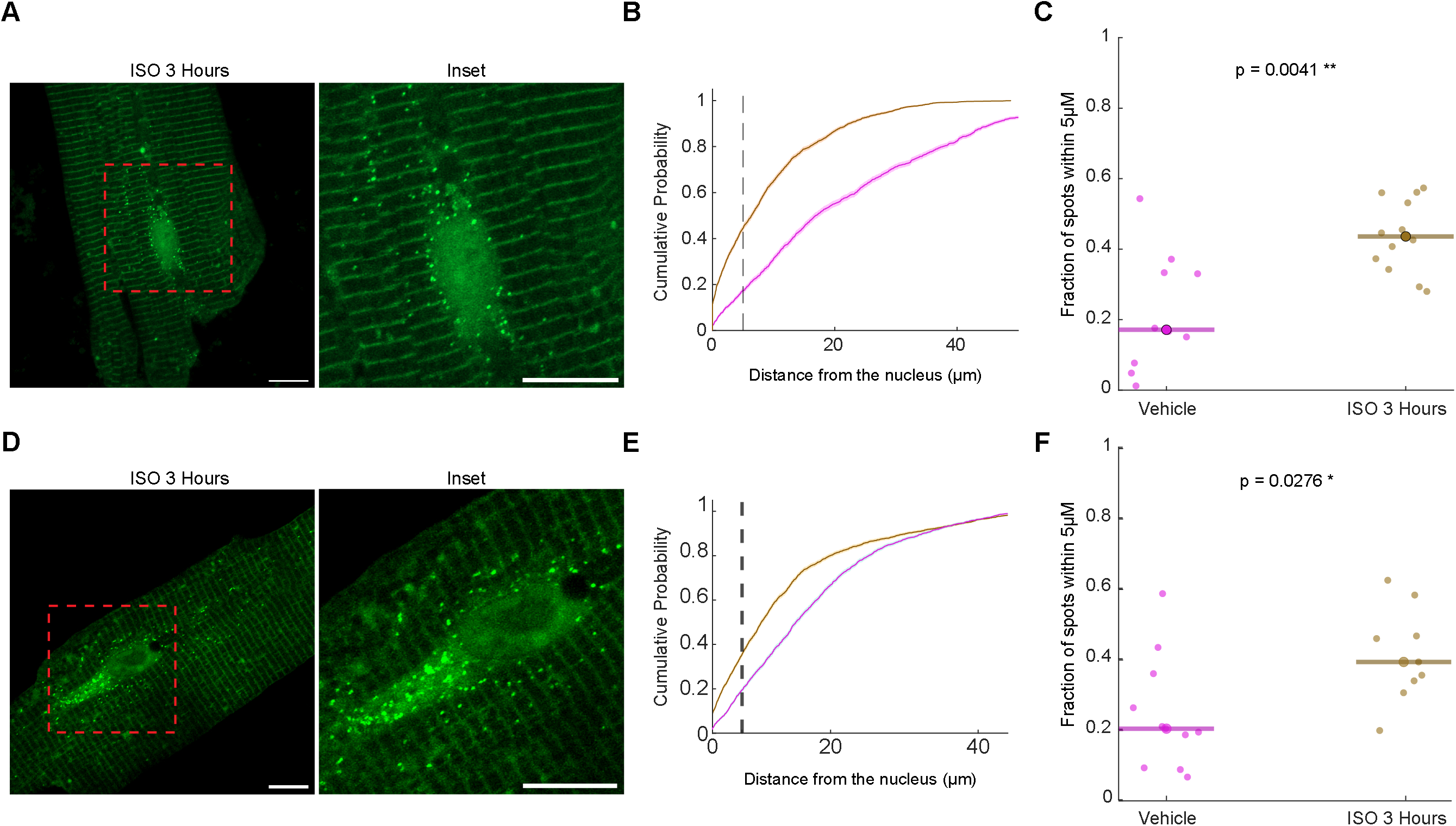
β-adrenergic stimulation results in peri-nuclear activation of translation. **A**, Representative image of adult cardiomyocyte treated for 3 hours with the non-selective β-adrenergic agonist isoprenaline (ISO). Red dashed square magnifies the perinuclear region, revealing the accumulation of active translation sites. **B**, Cumulative distribution of active translation sites as a function of distance from nuclear edge (vehicle – pink, isoprenaline - brown). **C**, Analysis of cumulative probability values at 5 μm distance from the nucleus for each experimental condition (vehicle: N=2 independent experiments, n=10 cells; isoprenaline: N=2 independent experiments, n=12 cells). Statistical significance was determined using Mann-Whitney U test (p = 0.0041). **D-F**, Same analyses as in A-C with cardiomyocytes transduced with SINAPs reporter that includes the 3’UTR of desmoplakin (*Dsp*), demonstrating that perinuclear accumulation does not depend on the 3’UTR. (vehicle: N=1 independent experiments, n=11 cells; isoprenaline: N=1 independent experiments, n=9 cells). Statistical significance was determined using Mann-Whitney U test (p = 0.0276).

**Supplemental Vido 1.** A 2:24 minute video is showing an adult cardiomyocyte transduced with the SINAPs reporter system treated with the translation inhibitor puromycin, resulting in disappearance of the bright translation spots, representing active translation sites.

**Supplemental Vido 2.** A 5 minute video showing an adult cardiomyocyte transduced with the SINAPs reporter system (green) and co-staining of microtubules (purple). A mobile active translation site moving along a microtubule in the long axis of the cardiomyocyte is highlighted with a yellow box, and a confined translation site that is mostly localized to the Z-line is highlighted with a red box.

## SUPPLEMENTARY METHODS

### Experimental Models and Animal Procedures Ethical Considerations

All animal procedures were reviewed and approved by the Technion Institutional Animal Care and Use Committee (IACUC) and were performed in strict accordance with institutional and national guidelines for the ethical use of animals in research. Every effort was made to minimize animal suffering and reduce the number of animals used in the study.

### Adult Rat Cardiomyocyte Isolation

Primary adult rat cardiomyocytes were isolated following established protocols.^1^ Adult Wistar rats were deeply anesthetized with ketamine and xylazine, and hearts were rapidly excised and immersed in ice-cold Krebs-Henseleit buffer containing 1 mM Ca²⁺ (KHB-Ca²⁺). After aortic cannulation, hearts underwent retrograde Langendorff perfusion with calcium-free KHB for 5 minutes at 37°C. Enzymatic digestion proceeded with recirculating perfusion of Collagenase Type II solution (320 U/ml; Worthington Biochemical) in KHB for 15 minutes, followed by gradual calcium reintroduction to 0.75 mM for an additional 15 minutes.

Post-perfusion, ventricular tissue was mechanically dissociated in KHB-Ca²⁺ enzyme solution through two sequential 5-minute gentle trituration steps. Myocytes were purified via centrifugation (70×g, 2 minutes) and resuspended in 2% BSA, 1 mM CaCl₂-KHB solution. Calcium concentration was incrementally restored to physiological levels (1.75 mM) through three sequential CaCl₂ additions over 15 minutes. The resulting viable myocytes were pelleted via centrifugation, resuspended in DMEM supplemented with 5% fetal calf serum, and seeded onto Matrigel-coated (Corning, #354234) µ-Dish 35 mm plates (Ibidi, #80416).

### Molecular Biology and Genetic Tools SINAPS Reporter System Design

We used the single-molecule imaging of nascent peptides (SINAPS) reporter system to visualize active translation in cardiomyocytes.² The reporter construct comprised 24 SunTag peptide domains, a 240-amino acid open reading frame based on oxygen-insensitive blue fluorescent protein (oxBFP), and a C-terminal auxin-induced degron (AID) sequence. To simultaneously track both mRNA and active translation, some constructs also contained 24XMS2-stem loops (v5) at their 3’UTR.

For alternative reporters, we incorporated various sequences into the 3’UTR of our reporter. These included the non-sarcomeric beta-actin (*Actb1*), as well as either the 3’UTR of sarcomeric actin (*Actc1*) or its complete sequence including the 5’ and 3’UTR and coding sequence (*Actc1*).

Additionally, we incorporated the 3’UTR of desmoplakin (*Dsp*). The pUbC-FLAG-24xSuntagV4-oxEBFP-AID-baUTR1-24xMS2V5-Wpre (Addgene plasmid #84561; http://n2t.net/addgene:84561; RRID) and the pUbC-OsTIR1-myc-IRES-scFv-sfGFP (Addgene plasmid # 84563; http://n2t.net/addgene:84563; RRID:Addgene_84563) were a gift from Robert Singer and Bin Wu.

### Adenoviral Vector Engineering

Custom adenoviral constructs encoding the SINAPs reporter and components were generated using Gateway cloning technology (Thermo Fisher). Target coding sequences were PCR-amplified from an NRVM cDNA library with high-fidelity polymerase and cloned into Gateway’Entry’ plasmids. After sequence verification by Sanger sequencing, validated inserts were subcloned into pAd/CMV/V5-DEST adenoviral expression vectors (Thermo Fisher) through LR recombination.

### Viral Production and Purification

Replication-incompetent adenoviral particles were produced in HEK293 cells following established protocols.^3^ Briefly, HEK293 cells at 80% confluency were transfected with PacI-linearized adenoviral plasmids using Polyethylenimine (PEI). Viral particles were harvested 7-10 days post-transfection, amplified through sequential passages, and purified using Iodixanol gradient ultracentrifugation.

### Cardiomyocyte Transduction

Viral transduction was performed 2-hours after plating by incubating cardiomyocytes with adenoviral vectors for 4 hours, followed by replacement with serum-free DMEM supplemented with ITSx100 (Gibco, #51500056) and Cytochalasin D (Cayman, #11330). To induce degradation of completed reporter proteins prior to experimentation, culture medium was exchanged with medium containing Indole-3-acetic acid (IAA) (Sigma-Aldrich, #I3750) 24 hours before imaging, followed by overnight incubation.^4^ This treatment ensured that observed fluorescent signals represented nascent peptides at translation sites rather than fully synthesized proteins.

### Pharmacological Treatments Translation Validation Assays

To confirm the specificity of our translation visualization system, we employed the translation elongation inhibitor Puromycin (Goldbio, #P-600) at a final concentration of 100 µg/mL as described.^2^

### Microtubule Network Manipulation

For microtubule depolymerization experiments, culture medium was replaced with medium containing 10 µM Nocodazole (Sigma-Aldrich, #M1404). Following a 24-hour incubation period, cells were subjected to live-cell microscopy to examine the effects of microtubule disruption on translation site distribution. Control cells were treated with an equivalent volume of vehicle (DMSO).

### Adrenergic Signaling Modulation

To investigate adrenergic regulation of mRNA translation, cardiomyocytes were exposed to adrenergic stimulation using one of two approaches:

1. ***α*-adrenergic stimulation**: Cells were treated with 50 µM Phenylephrine (Sigma-Aldrich) in 0.2% ascorbic acid (Sigma-Aldrich) dissolved in PBS. This treatment was applied for varying durations (0, 1, 2, 3, 6, 12, 24, or 72 hours) to assess time-dependent effects on translation site distribution. For time-lapse experiments, individual cells were continuously monitored for up to 6 hours after treatment initiation.
2. ***β*-adrenergic stimulation**: Cells were treated with 1 µM Isoprenaline (Sigma-Aldrich) dissolved in ultrapure water for 3 hours.

### mTOR Pathway Inhibition

To examine the role of mTOR signaling in adrenergic-induced translation site redistribution, the culture medium was replaced with medium containing 50 nM Torin1 (Tocris, #4247), a potent ATP-competitive inhibitor of mTOR kinase activity that blocks both mTORC1 and mTORC2 complexes.^5^ Cells were pre-treated with Torin1 for 10 minutes before exposure to phenylephrine (50 µM).

### ERK Inhibition

To examine the role of ERK signaling in adrenergic-induced translation site redistribution, the culture medium was replaced with medium containing 10 µM U0126 (Sigma-Aldrich, U120), a specific inhibitor of MEK1 and MEK2. Cells were pre-treated with U0126 for 10 minutes before exposure to phenylephrine (50 µM).

## Imaging Methods

### Live-Cell Microscopy

High-resolution imaging was performed using a Zeiss LSM 900 with Airyscan2 super-resolution system coupled to an Axio Observer 7 inverted microscope equipped with a PECON incubation insert for precise regulation of temperature (37°C), CO₂ (5%), and humidity. Image acquisition employed a Plan-Apochromat 63×/1.4 NA Oil DIC M27 objective (#420782-9900-799, Zeiss) with solid-state lasers at 488 nm and 640 nm.

For time-lapse imaging, cells were maintained in the incubation chamber with stable environmental conditions, and images were acquired at defined intervals (0, 1, 2, 3, and 6 hours) to monitor the redistribution of translation sites following treatment with vehicle or phenylephrine. To minimize photobleaching during extended imaging sessions, laser power was reduced to the minimum required for adequate signal detection, and the number of z-slices was optimized to capture the full cell volume while minimizing exposure.

To track both translation sites (TLS) and the microtubular network simultaneously, we used Tubulin Tracker™ (Invitrogen™, T34077) at a final concentration of 1 µM according to the manufacturer instructions. Cells were incubated with the Tubulin Tracker for 60 minutes before imaging and maintained in imaging medium containing the tracker during acquisition to allow for continuous visualization of the microtubule network alongside translation sites.

To track both the mRNA of the reporter and its translation site (TLS) we transduced the cells with three Adeno-viral vectors: Ad-scFV-sfGFP-OsTIR; Ad-MCP-2xNLS-HaloTag; Ad-SIN-null. Right before the live imaging session, we washed the cells with serum-free DMEM and then incubated cells with serum-free DMEM supplemented also with 200nM Janelia Fluor 635 (JF-635) ligand for 15 minutes to label the MCP-Halotag proteins that are getting bound to the MS2-loops on the mRNA of our reporter, enabling us to live image both mRNA and its live translation.

### Fluorescence Recovery After Photobleaching (FRAP)

For FRAP experiments, a region containing a single translation site was targeted using the 488 nm laser within a precisely defined region of interest. The target region was then photobleached with 5 iterations of 100% laser power (488 nm) to ensure complete inactivation of fluorophores. Post-bleach recovery was monitored by acquiring images every 20 seconds for 10 minutes.

When the translation site was bleached after the pre-images, we used the position of the TLS determined in the pre-bleaching images to calculate intensity until the TLS reappeared. Afterwards, we used the newly emerged TLS position to calculate intensity. The recovery curves were normalized with the integrated intensity value of the TLS measured in the pre-bleaching images. A nonlinear least square fit (lsqcurvefit in Matlab) was used to fit the theoretical curve to extract translation kinetic parameters.

For each experiment, unbleached control translation sites were monitored in the same field of view to account for potential photobleaching during acquisition and confirm that signal recovery in the target region was due to new protein synthesis rather than fluorophore diffusion or other artifacts.

### Immunofluorescence and Confocal Microscopy

For immunofluorescence analysis, cardiomyocytes were fixed with freshly prepared 4% paraformaldehyde in PBS for 15 minutes at room temperature, permeabilized with 0.2% Triton X-100 for 10 minutes, and blocked with 5% normal goat serum in PBS for 1 hour. Primary antibodies were diluted in blocking solution and applied overnight at 4°C with gentle agitation. The following primary antibody was used: anti-α-actinin-2 (ACTN2) (1:500, clone EA-53, Sigma-Aldrich) to visualize sarcomeric Z-lines. After extensive washing (3 × 10 minutes in PBS), cells were incubated with appropriate Alexa Fluor-conjugated secondary antibodies (1:500, Invitrogen) for 1 hour at room temperature. Nuclei were counterstained with DAPI (1:1000, Sigma-Aldrich).

High-resolution imaging was performed using a Zeiss LSM 900 confocal microscope with Airyscan2 detector. Z-stacks were acquired with optimal sampling parameters according to Nyquist criteria (XY: 0.05 µm, Z: 0.15 µm steps). Raw Airyscan images were processed using ZEN Blue software (Zeiss) with default settings to achieve resolution enhancement while maintaining signal-to-noise integrity. For quantitative comparative analyses, identical acquisition settings were rigorously maintained across all experimental conditions.

Line scan analysis was performed to examine the co-localization of scFv-sfGFP signal with α-actinin-positive Z-lines.

### Single-Molecule Fluorescence In Situ Hybridization (smFISH)

For visualization of specific RNA populations, we employed smFISH using Stellaris RNA FISH probes (Biosearch Technologies). For 18S rRNA detection, probes were labeled with TAMRA fluorophores (yellow). For poly(A) mRNA detection, probes targeting the poly(A) tail were labeled with Quasar-670 fluorophores (red). Probe sets were designed to target specific regions of interest within the transcripts while avoiding potential cross-hybridization with other cellular RNAs.

smFISH was performed according to the manufacturer’s optimized protocol and as previously described.^6^ Briefly, cells were fixed with 4% paraformaldehyde, permeabilized, and hybridized with probe sets (50 nM final concentration) in hybridization buffer overnight at 37°C in a humidified chamber. Following stringent washing steps to remove unbound probes, samples were counterstained with DAPI and mounted in anti-fade medium.

## Data Analysis

### Translation Site Tracking and Mobility Classification

Comprehensive analysis of translation site dynamics was performed using a custom analytical pipeline integrating existing software packages and novel algorithms implemented in MATLAB (MathWorks). Initial particle detection and tracking utilized the ImageJ plugin TrackMate.^7^

Translation sites were classified based on their mobility profiles using a histogram-based approach that analyzed the statistical distribution of displacements from the center position. Key classification parameters included the 90th percentile of distances from movement center and maximum excursion distance. Based on these metrics, translation sites were categorized as either “confined” (remaining proximal to their center of movement with limited maximum displacement) or “mobile” (exhibiting frequent excursions from center or larger maximum displacements).

For visualization of translation site trajectories, displacement vectors were color-coded according to time progression throughout the imaging interval, with axes showing the relative position of translation sites with respect to the adjacent Z-line.

### Spatial Relationship Analysis

To elucidate the spatial relationship between translation sites and sarcomeric architecture, we developed a computational framework to characterize translation site dynamics relative to Z-lines. For each translation site, we identified the nearest Z-line and established a transformed coordinate system where the Z-line was positioned at Y=0 through appropriate rotation and translation transformations. This approach enabled intuitive visualization and quantitative analysis of movement patterns relative to the underlying sarcomeric infrastructure. The relationship between mobility classification and Z-line proximity was assessed by comparing mean distances from Z-lines between confined and mobile translation populations.

For comparing TLS distributions between different protein groups, the distance from each TLS to the nearest Z-line was calculated using a Euclidean distance transform of the Z-line mask. Only TLS points located within the cell boundary mask were included in the analysis. Then, we employed Jensen-Shannon Divergence (JSD) to measure distance between probability distributions. Statistical significance of JSD values was determined through permutation testing (1000 permutations).

Further analysis examined the association of translation sites with cytoskeletal elements. Translation sites were categorized based on their proximity to microtubules, Z-lines, or the junctions between these structures. For analysis of directional preference, translation sites were categorized by mobility using a 200 nm displacement threshold to identify mobile structures. Angular measurements relative to the cellular longitudinal axis (0°) and transverse axis (90°) were calculated for each translation trajectory. Directional preferences were statistically compared between the complete translation site population and the mobile subset to determine whether mobility characteristics influenced directional behaviors.

### Active Translation Sites Distance Quantification and Statistical Analysis

To quantify the spatial distribution of active translation sites (TLS) relative to the nucleus, we developed an image-based analysis pipeline. For each image, we created a binary nuclear mask and computed a Euclidean distance map to measure the minimum distance from each identified TLS spot to the nuclear boundary.

To analyze active translation sites (TLS) localization patterns across different timepoints following treatment, we employed cumulative distribution functions (CDFs) to compare the complete spatial distribution of TLS spots relative to the nucleus. The CDF approach allowed us to visualize and quantify the progressive changes in TLS nuclear proximity across the entire range of measured distances rather than using arbitrary thresholds.

For statistical comparisons, we calculated the fraction of spots within 5 μm of the nuclear boundary for each image across all conditions, as this distance represents the perinuclear region. For comparisons between multiple treatment groups, statistical significance was determined using the Benjamini-Hochberg false discovery rate (FDR) correction for multiple comparisons. For direct comparisons between two conditions, statistical significance was determined using the Mann-Whitney U test.

For probability distribution function (PDF) analysis of translation site localization in control versus nocodazole-treated cells, we calculated the normalized frequency of translation sites at different distances from the nucleus and plotted the distribution to visualize shifts in localization patterns.

